# PAX proteins *in silico* prediction in *Lissachatina fulica*

**DOI:** 10.1101/2024.07.16.603807

**Authors:** Irina N. Dominova, Kristina Golovneva, Nadezhda Korshunova, Valery V. Zhukov

## Abstract

Prediction of snail PAX group proteins in *Lisachatina fulica* was performed. For this this purpose, homologous sequences to already known PAX proteins of *Cephalopoda, Gastropoda* and *Bivalvia* molluscs were predicted. The presence of certain domains (Paired box, Homeodomain and octapeptide) in the structure of the primary predicted sequences of *Lisachatina fulica* proteins from the GigaDB was analyzed. Also, to confirm that the sequences found belong to the PAX family proteins, their secondary and 3D structures were predicted, and their DNA-binding capacity was compared to the control Paired box domain using molecular docking. As a result, 8 proteins of *Lisachatina fulica* containing PAX characteristic sequences were identified. Based on the results of domain structure search and phylogenetic analysis, the proteins were categorized into 5 subfamilies of PAX proteins, namely I, III, IV, β, and Pox-Neuro. The result provides a starting point for the search for the functional role of PAX proteins in the *Lisachatina fulica*.

## Introduction

Paired-box (*PAX*) genes encode a family of highly conserved transcription factors found in both vertebrates and invertebrates. PAX proteins get their name from the “paired-box,” a region of homology firstly described as located between the paired (Prd) and gooseberry (Gsb) protein’s genes of *Drosophila*. Later this motif was found to encode sequence-specific DNA-binding activity (Underhill 2012). PAX proteins are defined by the presence of a highly conserved paired domain. This DNA-binding domain is 128 amino acids long and called a bipartite paired domain. It, consists of an N-terminal subdomain (PAI) and a C-terminal subdomain (RED) connected by a linker. These subdomains form a three-helix fold, with the C-terminus including of a helix-turn-helix (HTH) motif that binds DNA. In addition, the PAI subdomain includes an N-terminal beta-turn and a beta-hairpin, called a “wing,” also involved in DNA binding (Underhill 2000; Apuzzo et al. 2004). As well, some PAX proteins contain a 60 amino acid helix-turn-helix homeodomain and an octapeptide motif distal to the Paired-box domain (Panneerselvam et al. 2019). A total of 9 variants of PAX proteins have been found in the animal. These proteins are grouped into 4 common subfamilies based on their structure and conserved functions:

1. subfamily I (PAX1 and PAX9) contains Paired box domain and octapeptide,
2. subfamily II (PAX2, PAX5 and PAX8) – Paired box domain, partial homeodomain and octapeptide. However, Navet et al. reported that Pax2/5/8 proteins in *Lophotrochozoa* lack a homeodomain (Navet et al. 2017),
3. subfamily III (PAX3 and PAX7) – Paired box domain, complete homeodomain and octapeptide. Meanwhile, it should be noted that *Gastropoda* lacks the octapeptide (Navet et al.2017),
4. subfamily IV (PAX4 and PAX6) – Paired box domain and complete homeodomain (Mansouri et al. 1999). Also, two additional subfamilies were found in the invertebrates:
5. subfamily Pox-Neuro contains a Paired box domain and octapeptide (Friedrich 2015), however, the latter is absent in *Lophotrochozoa*. Meanwhile, the motif [VI]PGLSYP[KR][IL]V is found after the paired domain in *Mollusca* (Navet et al. 2017),
6. subfamily PAX-β is declared to be specific to the *Lophotrochozoa* (Schmerer et al. 2009), containg only the Paired box domain (Franke et al. 2015).

Thus, all PAX proteins contain a similar highly conserved DNA-binding motif; wherein, each of them regulates the transcription of various genes (Thompson et al. 2021). For instance, several studies have found that PAX family proteins play important roles in the development of various organs and tissues, including the thymus (PAX1 and PAX7), vertebrae (PAX1), ear (PAX2 and PAX8), kidney (PAX2), central nervous system (CNS) (PAX2, PAX5, PAX8, PAX6, PAX3 and PAX7), cardiovascular system, melanocytes, Schwann cells (PAX3 and PAX7), pancreas (PAX4 and PAX6), B lymphocytes (PAX5), eye (PAX6), skeletal muscles (PAX3 and PAX7), thyroid (PAX8), and teeth (PAX7 and PAX9) (Blake and Ziman 2014), also Pox-Neuro is involved in the CNS and peripheral nervous system development (Jiang et al. 2015). The role of PAX-β is not completely known. However, it has been found in *Helobdella austinensis*, where it expresses at early embryogenesis and appears to be involved in the transition from spiral to symmetrical segmentation (Schmerer et al. 2013).

At the moment structure and function of many PAX proteins of vertebrates are quite well studied, that opposite to invertebrates, particularly *Mollusca*. However, it is known that some PAX proteins of molluscs perform similar functions to that in vertebrates. For instance, it has been shown that PAX2, PAX5, and PAX8 proteins are involved in squid brain regionalization and the development of CNS. These proteins are also expressed during sensory systems development, including the eyes of some cephalopod and gastropod molluscs (Wollesen et al. 2015; Scherholz et al. 2017; Janeschik et al. 2022). For example, *Haliotis asinina*, PAX2, PAX5, and PAX8 proteins are expressed in statocysts (O’Brien and Degnan 2003). As well, expression of PAX family proteins are found in regions of sensory cells developing, that argues their involvement in its ontogenesis. Moreover, in molluscs these proteins are expressed only in ectoderm derivatives, suggesting their role in differentiation of this tissue (Wollesen et al. 2015). Wherein, in *Sepia officinalis* there is lack of PAX6 expression in developing retina, while this one is commonly found in other species. However, in other organs, such as skin and gills, the expression of PAX2, PAX5, and PAX8 is ever observed (Navet et al. 2017).

In molluscs, PAX6 protein, similar to vertebrates, controls the eye development. Vertebrates have four isoforms of PAX6 protein that control specific groups of genes, while insects have duplicated Pax6 genes for eye development. Yoshida et al. reported that in squid, five PAX6 isoforms, formed by alternative splicing, are responsible for eye development, resembling the mechanism in vertebrates. However, duplicated gene forms typical for insects are absent (Yoshida et al. 2014).

The study of mollusc PAX family proteins in molluscs should consider the morphological diversity within the phylum *Mollusca*, which include seven classes according to the Taxonomy Browser (https://www.ncbi.nlm.nih.gov/Taxonomy/Browser/ (accessed on 03 April 2023)). Special attention should be paid to the abilities of certain molluscs to regenerate individual body parts (Imperadore et al. 2017). This ability is well developed in cephalopod molluscs. In addition to octopuses, the land snail *Lissachatina fulica* (Férussac, 1821) is capable of regenerating specific body parts, such as the eye tentacle along with the eye. This regeneration ability was first demonstrated about 20 years ago (Bobkova et al. 2004). However, the available data on eye regeneration in this snail include only microscopic and electrophysiological examinations, as well as behavioral tests. The molecular genetic component has not been explored (Tuchina and Meyer-Rochow 2010). Considering the crucial role of PAX family proteins in eye development and formation, we believe that they likely play a significant part in the regeneration process. Identifying the molecular and genetic components involved in the regeneration mechanism is challenging due to the lack of an annotated genome for *L. fulica*. Therefore, it is essential to predict the amino acid sequences of PAX family proteins in the *L. fulica* through bioinformatic analysis. This will help shed light on their potential involvement in the morphogenesis and regeneration mechanisms of this gastropod mollusc.

Thus, to search for and conduct bioinformatics analysis of amino acid sequences containing the Paired box domain in *L. fulica*, it is necessary to: 1) predict sequences homologous to already known PAX proteins of molluscs; 2) confirm the presence of Paired box domain sequences and other functional elements in the structure; 3) predict the secondary and 3D structures of the found Paired box-containing sequences; 4) use molecular docking to compare the DNA-binding ability of the found sequences with the control Paired box domain; 5) use phylogenetic and BLAST analyses to determine whether the found Paired box-containing sequences belong to certain subfamilies of PAX proteins.

## Materials and Methods

### Compiling data from databases

For the PAX protein prediction in *Lissachatina fulica* (Férussac, 1821), we selected three mollusc classes: *Cephalopoda, Gastropoda* and *Bivalvia*. PAX proteins and Paired box domain-containing proteins from these three mollusc classes were collected from the NCBI Protein database (https://www.ncbi.nlm.nih.gov/protein/ (accessed on 31 October 2022)) using BLAST search with two algorithms BLASTp and PSI-BLAST against to the PF00292 domain (https://www.ebi.ac.uk/interpro/entry/pfam/PF00292/ (accessed on 31 October 2022)). Performing protein BLAST (BLASTp) and psiBLAST searches allows the identification of sequences that are absent in phylogenomic databases (Duek et al. 2021). We excluded proteins with PREDICTED and LOW QUALITY status, as well as hypothetical protein, from further analysis. Thus, 91 sequences of PAX and Paired box domain-containing proteins (Table S1) were collected for further analysis.

The primary predicted sequences of *L. fulica* proteins were taken from Guo’s article (Guo et al. 2019) supplementary materials deposited in the GigaDB database (http://gigadb.org/dataset/100647 (accessed on 31 October 2022)).

### Protein prediction

Computationally identifying PAX-containing proteins in the *L. fulica* was performed using the HMMER software (version 3.4) (http://hmmer.org/ (accessed on 31 October 2022)) from National Human Genome Research Institute, USA (Mistry et al. 2013). To accomplish this, a multiple sequence alignment of the 91 previously collected PAX-containing molluscan proteins was conducted using the Unipro UGENE software (version 45.1) from Unipro, Russia (Okonechnikov et al. 2012). Clustal Omega (Sievers and Higgins 2018) and MUSCLE (Edgar 2004) algorithms integrated in UGENE were used to align the amino acid sequences. Initial alignment was carried out by Clustal Omega with the following parameters: number of iterations — 100, max number of guide-tree iterations — 100, max number of HMM (Hidden Markov Models) iterations — 100. Subsequently, the alignment was refined using MUSCLE with a maximum of 100 iterations, using the entire amino acid sequences.

Based on the obtained alignment, an HMM3-profile was constructed in the Unipro UGENE. This profile was then used to predict the studied protein sequences in the *L. fulica* using the *hmmsearch* algorithm for protein sequence analysis, which is integrated into HMMER. This algorithm employs the HMM profile built based on the multiple sequence alignment query and searches for it in the target sequence database (Finn et al. 2011). In our case, the target sequence database consisted of the primary predicted *L. fulica* protein sequences deposited in the GigaDB database.

After prediction, sequences with an E-value lower than the set threshold of 10^−5^ were selected for further analysis and verification.

### Verification (Domain search, GO annotation, subcellular location)

Confirmation of Paired box domain presence in the identified *L. fulica* sequences was performed using the InterProScan website (https://www.ebi.ac.uk/interpro/ (accessed on 8 February 2023)). InterProScan allows for the integrative classification of protein sequences into families and the identification of functionally important domains and conserved regions (Jones et al. 2014). InterPro integrates 13 protein signature databases: CATH-Gene3D, the Conserved Domains Database (CDD), HAMAP, PANTHER, Pfam, PIRSF, PRINTS, SMART, the Structure-Function Linkage Database (SFLD), SUPERFAMILY and TIGRFAMs, making it the most comprehensive resource in the world for protein families, domains, and functional sites (Paysan-Lafosse et al. 2022).

Multiple sequence alignments were also conducted using the UGENE software, employing the Clustal Omega and MUSCLE algorithms with the same parameters as those used to create the HMM-profile. These alignments aimed to identify and specify the positions of conserved domains in the predicted sequences. As reference sequences for conserved domains used were smart00351 Paired Box domain and smart00389 Homeodomain consensus (http://smart.embl.de/ (accessed on 15 March 2023)), pfam00292 ‘Paired box’ domain and pfam00046 Homeobox consensus (https://www.ebi.ac.uk/interpro/entry/pfam/#table (accessed on 15 March 2023)). Additionally, the octapeptide and specific motif sequences were obtained from the articles by Navet et al. and Ziman and Kay (Navet et al. 2017; Ziman and Kay 1998).

### Secondary and 3D structure

The secondary structure prediction was performed using RaptorX-Property software (http://raptorx6.uchicago.edu/StructurePropertyPred/predict/ (accessed on 5 April 2023)) (Wang et al. 2016), which provides the percentage of regular (α-helixes, β-sheets) and irregular secondary structures (coils) for both the entire sequence and each individual amino acid.

The 3D structure of the predicted proteins was derived by AlphaFold2 Colab, employing MMseqs2 (v1.5.3) (https://colab.research.google.com/github/sokrypton/ColabFold/blob/main/AlphaFold2.ipynb (accessed on July 07, 2023)) (Mirdita et al. 2022). For each sequence, the algorithm generated five approximate 3D structures and ranked them according to the pLDDT reliability index, which ranges from 0 to 100. This index provides an estimation of the reliability of a specific region in the structure prediction. The higher the index value, the greater the accuracy of the predicted structure (Carugo 2023). To facilitate further analysis and visualization, only the 3D structure with the highest pLDDT index was selected for each sequence.

PyMOL (version 2.5.7) (https://pymol.org/2/#download/ (accessed on 30 November 2023)) was used to structure alignment, visualize the results, and prepare figures (Schrödinger and DeLano 2020).

### DNA-Protein docking simulation

DNA-protein docking was performed using high ambiguity driven protein-DNA docking (HADDOCK) web server version 2.4 (https://wenmr.science.uu.nl/haddock2.4/ (accessed on 29 November 2023)). DNA-protein interactions play a vital role in various biological processes, and it is crucial to comprehend their structural aspects to uncover their functional mechanisms. Molecular docking, a computational approach, is employed to forecast the binding between ligands and proteins. HADDOCK is a versatile software that combines information-based docking with explicit solvent refinement to produce precise models of DNA-protein complexes. By integrating these restraints, HADDOCK utilizes experimental data to guide the docking process and improve the accuracy of predicted complexes (Van Zundert et al. 2016).

The docking process utilized the 3D structures of the predicted amino acid sequences of PAX-containing *L. fulica* proteins, which were obtained using AlphaFold2 Colab as previously described, and a 3D structure of B-form DNA (PDB ID: 1BNA) (Drew et al. 1981). Additionally, the 3D structure of the Paired box domain (PDB ID: 1PDN) (Xu et al. 1995) with removed DNA was employed as control for the docking process. Only the Paired box domain sequence was considered for docking in the predicted *L. fulica* proteins. HADDOCK produces various clusters comprising diverse structural conformations of proteins obtained through water refinement of models. An analysis of the top 10 clusters was conducted, with the foremost cluster being deemed the most trustworthy according to the HADDOCK scoring system. The energy (Z)-score of this top cluster serves as a measure of its deviation from the average cluster, with a more negative score signifying a stronger affinity between the protein and DNA.

### Phylogenetic analysis

Phylogenetic analysis was conducted using the MEGA X software (version 10.2.6) (https://www.megasoftware.net/ (accessed on 01 July 2022)) (Kumar et al. 2018) through Maximum Likelihood and the Le and Gascuel (LG) model with frequency (+F) corrections (Le and Gascuel 2008), along with 1000 bootstrap replications (Felsenstein 1985). The evolutionary model was selected using the IQ-TREE Web Server (http://iqtree.cibiv.univie.ac.at/ (accessed on 02 August 2023)) (Trifinopoulos et al. 2016), which automatically determined the best-fit model for each alignment. The initial tree for the heuristic search was obtained automatically through Maximum Parsimony. A discrete gamma distribution (+G) with four categories was used to account for different evolutionary rates among sites. Additionally, a rate variation model (+I) was included in the analysis to allow for some sites to remain evolutionarily unchanged. The resulting phylogenetic tree was automatically selected based on the topology with the highest logarithmic likelihood value. The phylogenetic trees were visualized using FigTree software (version 1.4.4) (http://tree.bio.ed.ac.uk/software/figtree/ (accessed on 07 August 2023)) (Rambaut 2018). To construct the phylogenetic tree, in addition to the predicted *L. fulica* sequences, we also included sequences from five classes of *Mollusca* (*Gastropoda, Cephalopoda, Bivalvia, Aplacophora* and *Polyplacophora*). These classes were chosen to represent the diversity of PAX protein subfamilies.

### BLAST analysis

*L. fulica* predicted proteins were searched using BLASTp algorithm against gastropod, cephalopod, and bivalve PAX-containing proteins deposited in the NCBI GenBank’s protein database of Reference proteins (refseq_protein) (https://blast.ncbi.nlm.nih.gov/Blast.cgi (accessed on 12 July 2023)).

## Results

The analysis of PAX and PAX-containing protein sequences, selected from the BLAST search to build the HMM profile, revealed an unequal distribution among the mollusc classes. Specifically, *Gastropoda* representatives accounted for 29 sequences, *Cephalopoda* for 9 sequences, and *Bivalvia* for the majority with 53 sequences. Furthermore, the selected 91 protein sequences were also unevenly distributed among PAX groups or subfamilies. Group 1 was represented by only one PAX-1 protein, group 2 by 70 representatives (Pax-2: 21 proteins, Pax-5: 22 proteins, and Pax-8: 27 proteins), and group 4 by 20 Pax-6 proteins. Group 3 proteins (Pax-3 and Pax-7) was found to be absent in molluscan genomes.

### PAX-containing protein prediction and verification

The prediction of PAX-containing proteins using HMMER showed the presence of 10 such sequences (Table 1) in the *L. fulica*.

**Table 1.**
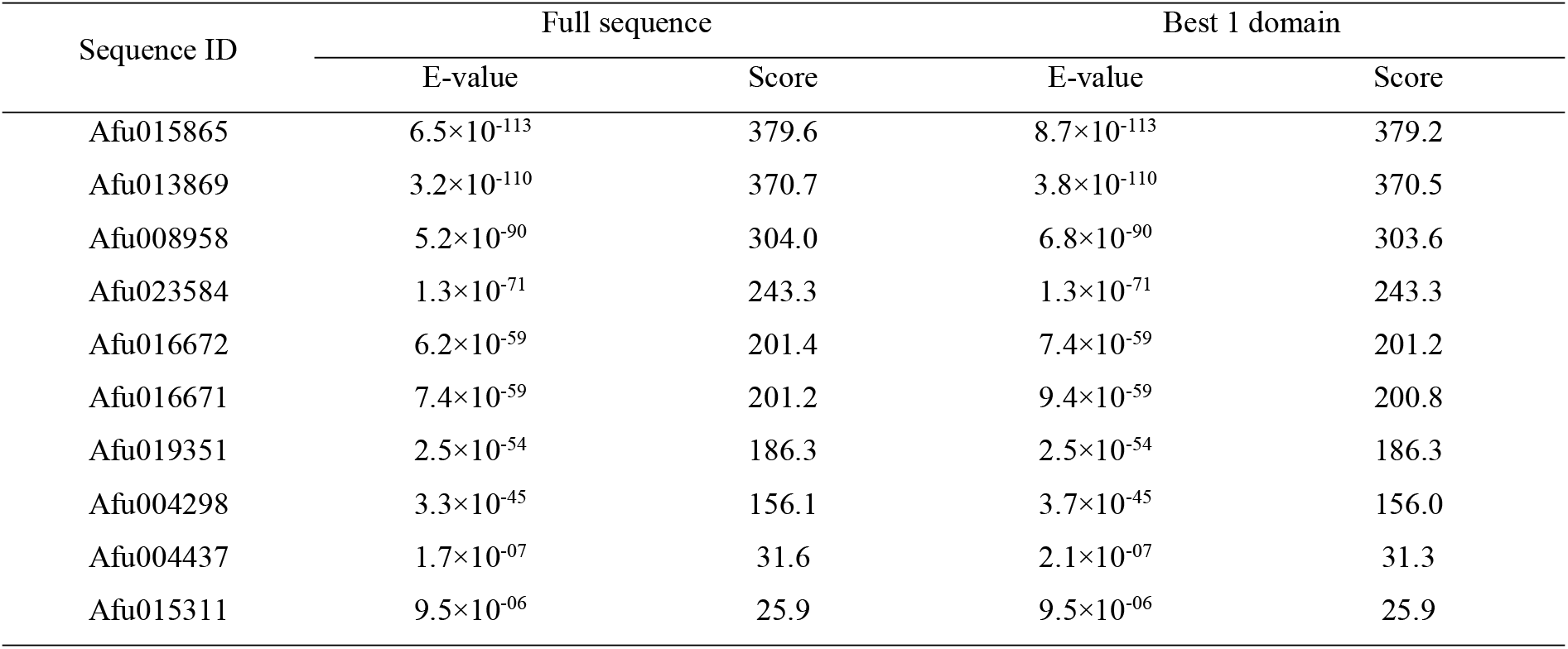
Significant results (E-value < 10^−5^) of the HMMER analysis for the prediction of PAX-containing proteins in *Lissachatina fulica*.

| Sequence ID | Full sequence |  | Best 1 domain |  |
| --- | --- | --- | --- | --- |
|  | E-value | Score | E-value | Score |
| Afu015865 | $6.5 \times 10^{-113}$ | 379.6 | $8.7 \times 10^{-113}$ | 379.2 |
| Afu013869 | $3.2 \times 10^{-110}$ | 370.7 | $3.8 \times 10^{-110}$ | 370.5 |
| Afu008958 | $5.2 \times 10^{-90}$ | 304.0 | $6.8 \times 10^{-90}$ | 303.6 |
| Afu023584 | $1.3 \times 10^{-71}$ | 243.3 | $1.3 \times 10^{-71}$ | 243.3 |
| Afu016672 | $6.2 \times 10^{-59}$ | 201.4 | $7.4 \times 10^{-59}$ | 201.2 |
| Afu016671 | $7.4 \times 10^{-59}$ | 201.2 | $9.4 \times 10^{-59}$ | 200.8 |
| Afu019351 | $2.5 \times 10^{-54}$ | 186.3 | $2.5 \times 10^{-54}$ | 186.3 |
| Afu004298 | $3.3 \times 10^{-45}$ | 156.1 | $3.7 \times 10^{-45}$ | 156.0 |
| Afu004437 | $1.7 \times 10^{-07}$ | 31.6 | $2.1 \times 10^{-07}$ | 31.3 |
| Afu015311 | $9.5 \times 10^{-06}$ | 25.9 | $9.5 \times 10^{-06}$ | 25.9 |

To verify whether the 10 predicted sequences belonged to PAX family proteins, a Paired box domain search was conducted using the InterPro web service (Table S2). As a result, two sequences (Afu015311 and Afu004437) were found not to belong to the PAX protein family because they only have a homeodomain without a Paired box domain. Therefore, the remaining 8 sequences were found to contain regions of varying lengths corresponding to the Paired box domain.

The presence of the Paired DNA-binding domain in the predicted sequences allows us to classify these proteins as transcription factors (GO:0006355, GO:0003677) based on GO categories. The subcellular location for all predicted sequences was determined to be the nucleus..

Multiple sequence alignments of the identified sequences with conserved Paired box domain (smart00351 and pfam00292), as well as secondary and 3D structures prediction (Fig. 1, Fig. S1), also confirmed the absence of the Paired box domain in Afu004437 and Afu015311 (not shown in Fig. 1). Additionally, a few sequences were found to contain partial Paired box domain. For example, Afu004298 contains only the first part of Paired box domain, specifically the PAI subdomain. Afu016672 and Afu016671 also contain a partial PAI subdomain because their sequences lack the first α-helix. Furthermore, Afu019351 lacks the second and part of the third α-helices of the PAI subdomain.

**Fig. 1.**
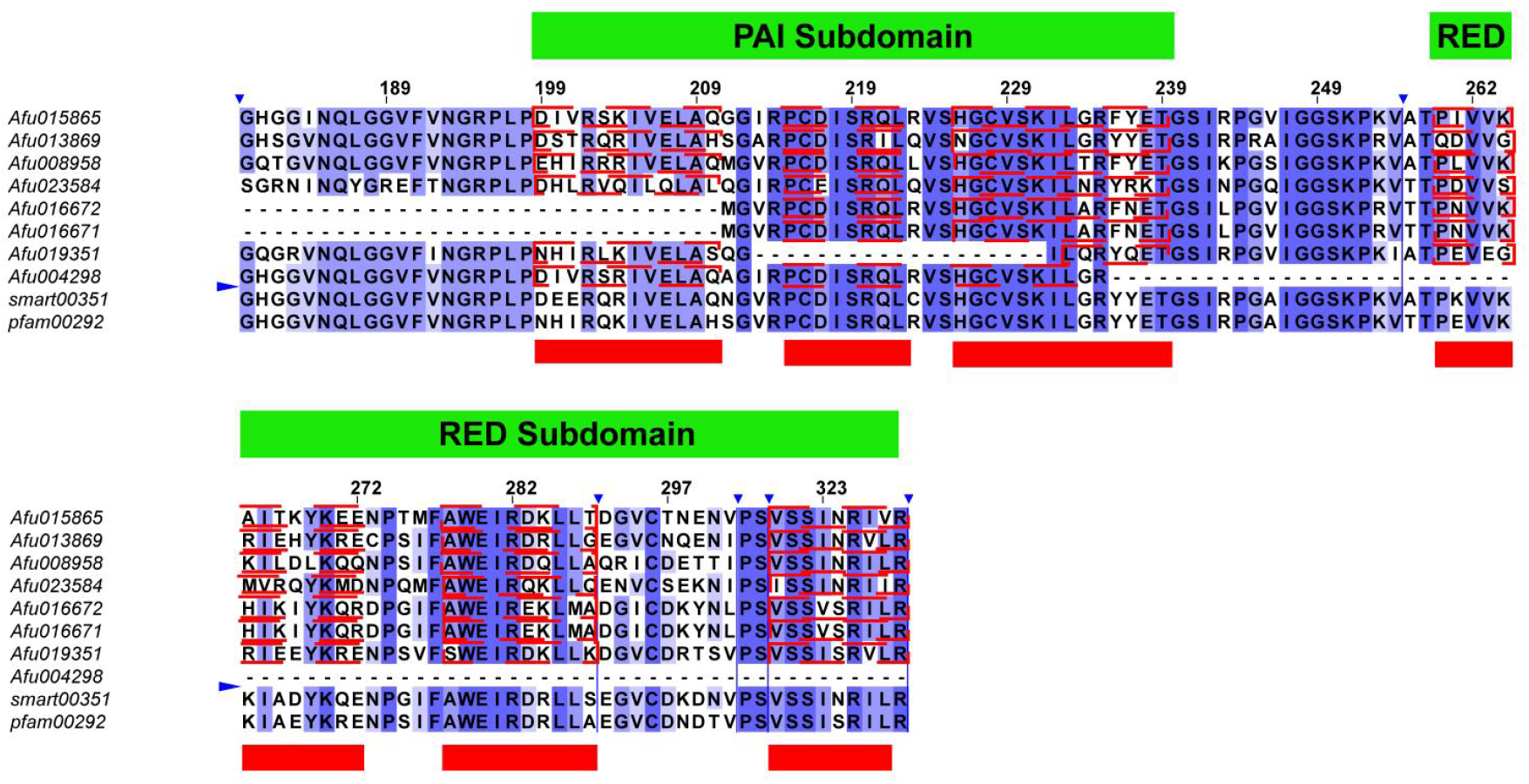
Alignment of amino acid sequences of *Lissachatina fulica* predicted proteins and Paired box domains (smart00351 and pfam00292). The most conserved regions are presented. Multiple sequence alignments were performed by the Clustal Omega algorithm and refined with the MUSCLE algorithm. Amino acid residues with >60% identity are highlighted in lilac colour. The red rectangles indicate the location of the α-helices in the conserved Paired box domains. The red lines represent the predicted position of the α-helices in the L. fulica amino acid sequences. Green rectangles indicate positions corresponding to the PAI and RED subdomains. The sign 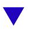 above the sequences indicates the excised regions. The single-letter amino acid designations correspond to the commonly used ones. The image was created using GNU Image Manipulation Program (GIMP) (version 2.10)

A similar alignment were performed for the Homeodomain (smart00389 and pfam00046) (Fig. 2). The results showed that the full Homeodomain is only present in Afu019351, Afu004437 and Afu015311. On the other hand, a partial Homeodomain, specifically its first α-helix, was detected in Afu008958 and Afu023584. All other sequences do not include Homeodomain.

**Fig. 2.**
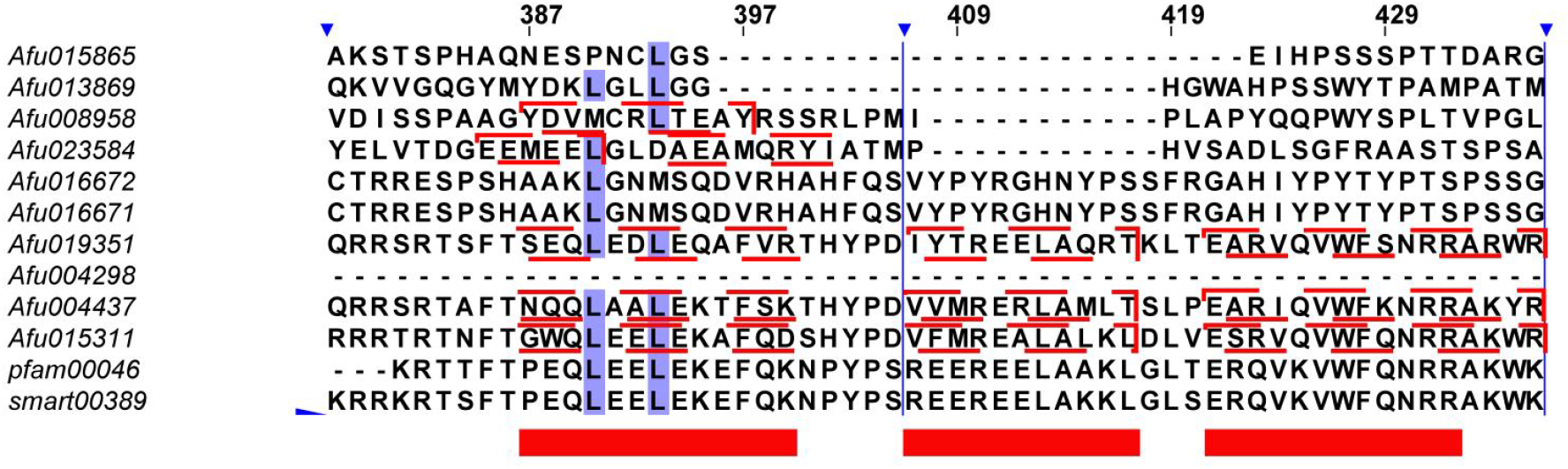
Alignment of amino acid sequences of *Lissachatina fulica* predicted proteins and Homeodomains (smart00389 and pfam00046). The most conserved regions are presented. Multiple sequence alignments were performed by the Clustal Omega algorithm and refined with the MUSCLE algorithm. Amino acid residues with >60% identity are highlighted in lilac colour. The red rectangles indicate the location of the α-helices in the conserved Homeodomains. The red lines represent the predicted position of the α-helices in the L. fulica amino acid sequences. The sign 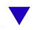 above the sequences indicates the excised regions. The single-letter amino acid designations correspond to the commonly used ones. The image was created using GNU Image Manipulation Program (GIMP) (version 2.10)

The search for octapeptide sequences revealed that the majority of the analyzed proteins contain them, with the exception of Afu019351, Afu004298, Afu004437, and Afu015311. Specifically, the sequence Afu015865 contains the octapeptide “DSELDLLT”, which on aligns most closely with octapeptides characteristic of PAX proteins of subfamilies I and III. On the other hand, the sequences Afu016671 and Afu016672 have the octapeptide “HSVHDILG,” differing by only one amino acid from the octapeptide specific to mouse and human PAX9 proteins (HSVTDILG). The octapeptide “YDKLGLLN” in sequence Afu013869 corresponds to octapeptides found in PAX subfamily II proteins. Among the amino acid residues of Afu008958, a “VPGLSYPRLV” sequence specific to Pox-Neuro proteins was identified. Additionally, the sequence “DKTLTLRRGY” was found in Afu023584, which is similar to the sequence found in Pax-beta proteins.

Furthermore, when analyzing the location of conserved domains, it is important to consider the size of the predicted sequences. The average length of paired-box-containing proteins in the *L. fulica* proteome is 1287 amino acids, with a maximum size of 3699 for Afu023584 and a minimum of 291 for Afu004298. The Paired box domain in PAX proteins is approximately 128 amino acids long, as mentioned earlier, and this condition is met in four identified sequences (Afu015865, Afu013869, Afu008958, and Afu023584) containing full PAI and RED subdomains.

### DNA-Protein docking findings

Docking was performed using HADDOCK 2.4 to analyze the interaction between predicted proteins of *L. fulica* and DNA (PDB ID: 1BNA), as well as the Paired box domain (PDB ID: 1PDN) and DNA (PDB ID: 1BNA) as a positive control (Table 2). The results showed that most of the predicted proteins had higher HADDOCK scores compared to the reference Paired-box domain (−82.8 ± 8.7), except for Afu004298 (−115.2 ± 18.3). However, when considering z-scores, Afu004298 (−2.3) exhibited the strongest DNA-binding ability, followed by Afu019351 (−1.9) and Afu008958 (−1.7), whereas Afu016671 (−1.1) had the lowest z-score in comparison with the Paired-box domain (−2.1). This suggests a potentially lower affinity of the predicted protein sequences of *L. fulica* (excluding Afu004298) for DNA compared to the reference Paired-box domain.

**Table 2.** Molecular docking simulation data for predicted sequences of *Lissachatina fulica* and Paired-box domain interacting with DNA represented by HADDOCK outputs.

| Characteristics | Afu015865 | Afu013869 | Afu008958 | Afu023584 | Afu016672 | Afu016671 | Afu019351 | Afu004298 | Paired-box domain (positive control) |
| --- | --- | --- | --- | --- | --- | --- | --- | --- | --- |
| HADDOCK score | $-48.9 \pm 3.5$ | $-5.0 \pm 6.7$ | $-28.0 \pm 15.0$ | $66.1 \pm 19.4$ | $-47.8 \pm 8.5$ | $-40.1 \pm 3.6$ | $-42.1 \pm 14.6$ | $-115.2 \pm 18.3$ | $-82.8 \pm 8.7$ |
| RMSD from the overall lowest-energy structure | $10.4 \pm 0.1$ | $4.8 \pm 0.5$ | $4.1 \pm 0.2$ | $4.5 \pm 0.3$ | $9.9 \pm 0.1$ | $8.0 \pm 0.3$ | $0.8 \pm 0.5$ | $1.8 \pm 1.4$ | $2.7 \pm 1.6$ |
| Van der Waals energy | $-70.4 \pm 14.4$ | $-60.0 \pm 7.0$ | $-55.0 \pm 3.6$ | $-65.0 \pm 13.2$ | $-57.2 \pm 5.2$ | $-57.4 \pm 2.6$ | $-91.7 \pm 1.8$ | $-62.9 \pm 14.3$ | $-53.7 \pm 11.2$ |
| Electrostatic energy | $-395.4 \pm 75.2$ | $-419.0 \pm 38.1$ | $-551.3 \pm 38.1$ | $-281.6 \pm 49.5$ | $-429.2 \pm 64.0$ | $-318.7 \pm 7.5$ | $-484.1 \pm 59.0$ | $-503.8 \pm 26.0$ | $-534.4 \pm 43.1$ |
| Desolvation energy | $24.3 \pm 3.6$ | $16.4 \pm 1.2$ | $22.2 \pm 4.6$ | $0.9 \pm 5.1$ | $16.1 \pm 1.2$ | $7.5 \pm 4.5$ | $24.7 \pm 4.2$ | $10.0 \pm 0.9$ | $23.8 \pm 0.9$ |
| Restraints violation energy | $762.8 \pm 113.2$ | $1224.3 \pm 44.0$ | $1150.9 \pm 95.6$ | $1865.3 \pm 117.7$ | $792.8 \pm 34.4$ | $736.0 \pm 25.5$ | $1217.1 \pm 74.1$ | $384.0 \pm 48.5$ | $539.5 \pm 50.9$ |
| Buried Surface Area | $2263.2 \pm 202.7$ | $2072.0 \pm 63.4$ | $1899.2 \pm 70.7$ | $1633.3 \pm 92.1$ | $1979.5 \pm 171.4$ | $1811.7 \pm 68.3$ | $2880.7 \pm 117.1$ | $1909.3 \pm 201.8$ | $2068.9 \pm 180.4$ |
| Z-Score | -1.3 | -1.4 | -1.7 | -1.4 | -1.6 | -1.1 | -1.9 | -2.3 | -2.1 |

Additionally, the predicted sequences showed higher Root Mean Square Deviation (RMSD) values, which measure the deviation of a molecular structure from the reference geometry. The RMSD of the reference Paired-box domain was 2.7 ± 1.6, while Afu019351 (RMSD = 0.8 ± 0.5) and Afu004298 (RMSD = 1.8 ± 1.4) had comparable or even smaller values among the predicted sequences. The highest RMSD value, indicating the worst deviation, was observed for Afu015865 (RMSD = 10.4 ± 0.1).

Similar trends with worse results were observed for Electrostatic energy, Restraints violation energy, and Buried Surface Area values when comparing the docking results of the Paired-box domain and predicted proteins with DNA. However, the Van der Waals energy and Desolvation energy values obtained for *L. fulica* sequences were lower than those obtained for the positive control. For example, most of the predicted proteins of *L. fulica* exhibited lower Buried Surface Area values, suggesting a smaller available surface area for DNA binding compared to the Paired-box domain, except for Afu019351 (2880.7 ± 117.1) and Afu015865 (2263.2 ± 202.7). The higher values of Electrostatic energy and Restraints violation energy for *L. fulica* sequences may also indicate a lower affinity of these proteins for DNA compared to the reference Paired-box domain sequence.

### Phylogenetic and BALST analyses

Analysis of the phylogenetic tree (Fig. 3), constructed using the amino acid sequences of PAX proteins of the five classes of Molluscs (*Gastropoda, Bivalvia, Cephalopoda, Aplacophora* and *Polyplacophora*), along with the predicted paired-box containing proteins of *L. fulica*, suggests that the sequences Afu015865, Afu004298, Afu016671, and Afu016671 belong to PAX subfamily I. Specifically, Afu015865 and Afu004298 form a common group with PAX-1 of *A. californica* (XP_012942774.1), while Afu016671 and Afu016671 cluster with PAX-1 of *O. bimaculoides* (XP_052829430.1). Afu016671 and Afu016671 also share a common clade with PAX-1 proteins from various *Gastropoda* representatives (*P. vulgata* and *A. califorica*), *C. gigas* and *O. bimaculoides*, as well as PAX-9 proteins (*P. ocellatus, E. marginata, M. yessoensis*).

**Fig. 3.**
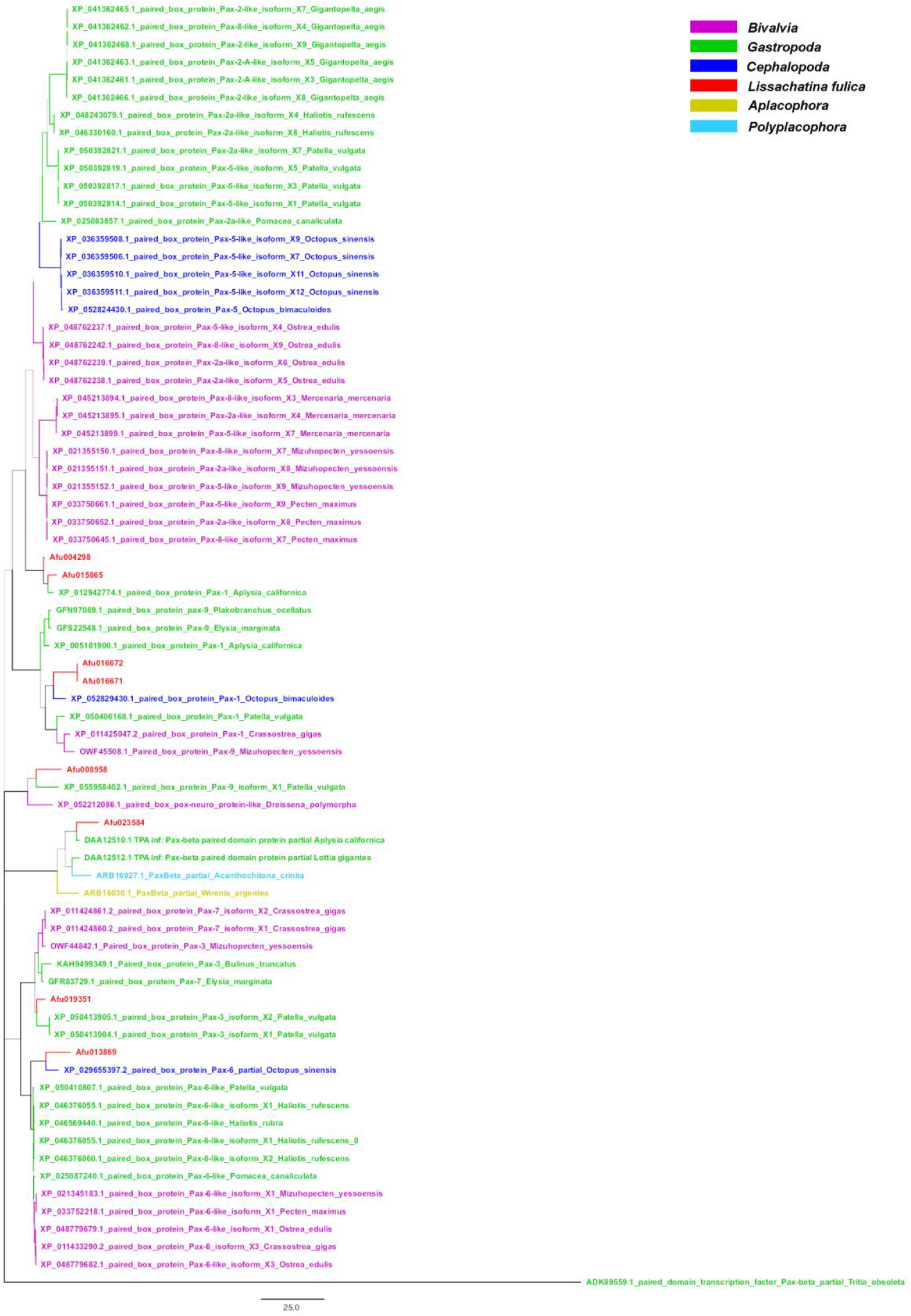
Phylogenetic tree of molluscan PAX and *Lissachatina fulica* predicted proteins. The tree was constructed for 73 amino acid sequences, with the final dataset containing a total of 1503 positions. The Maximum Likelihood (ML) statistical method and the LG correction model with 1000 bootstrap repeats were used. A discrete Gamma distribution was employed to model evolutionary rate differences among sites (four categories, +G, parameter = 0.3770). The rate variation model allowed for some sites to be evolutionarily invariable (+I, 0.13% sites). The tree is drawn to scale, and branch lengths represent the number of substitutions per site. The image was created using FigTree software (version 1.4.4) and GNU Image Manipulation Program (GIMP) (version 2.10)

The sequence Afu019351 belongs to a clade consisting of PAX subfamily III proteins from the classes *Gastropoda* (*P. vulgata, B. truncatus, E. margianta*) and *Bivalvia* (*C. gigas, M. yessoensis*). Although Afu013869 is a member of the cluster formed by PAX-6 proteins from representatives of three *Mollusca* classes (*Gastropoda, Bivalvia* and *Cephalopoda*), it is still somewhat distant from them. The closest protein to Afu013869 is PAX-6 of *O. sinesis* (XP_029655397.2).

The sequence Afu023584 not only belongs to the clade formed by PAX-beta proteins, but shares a common node with the PAX-beta protein of *A. californica* (DAA12510.1). On the other hand, the remaining analyzed sequence, Afu008958, is closest to the clade formed by two proteins belonging to different subfamilies of PAX proteins, specifically PAX-9 of *P. vulgata* (XP_055958402.1) and Pox-Neuro of *D. polymorpha* (XP_052212086.1). It is worth noting that only one amino acid sequence of Pox-Neuro is known for molluscs, so a more detailed phylogenetic analysis of Afu008958 is not possible.

In addition to phylogenetic analysis, separate BLAST searches were performed for *L. fulica* predicted proteins against *Gastropoda, Cephalopoda* and *Bivalvia* mollusc proteins to determine the percentage of identity of these amino acid sequences (Table 3).

**Table 3.**
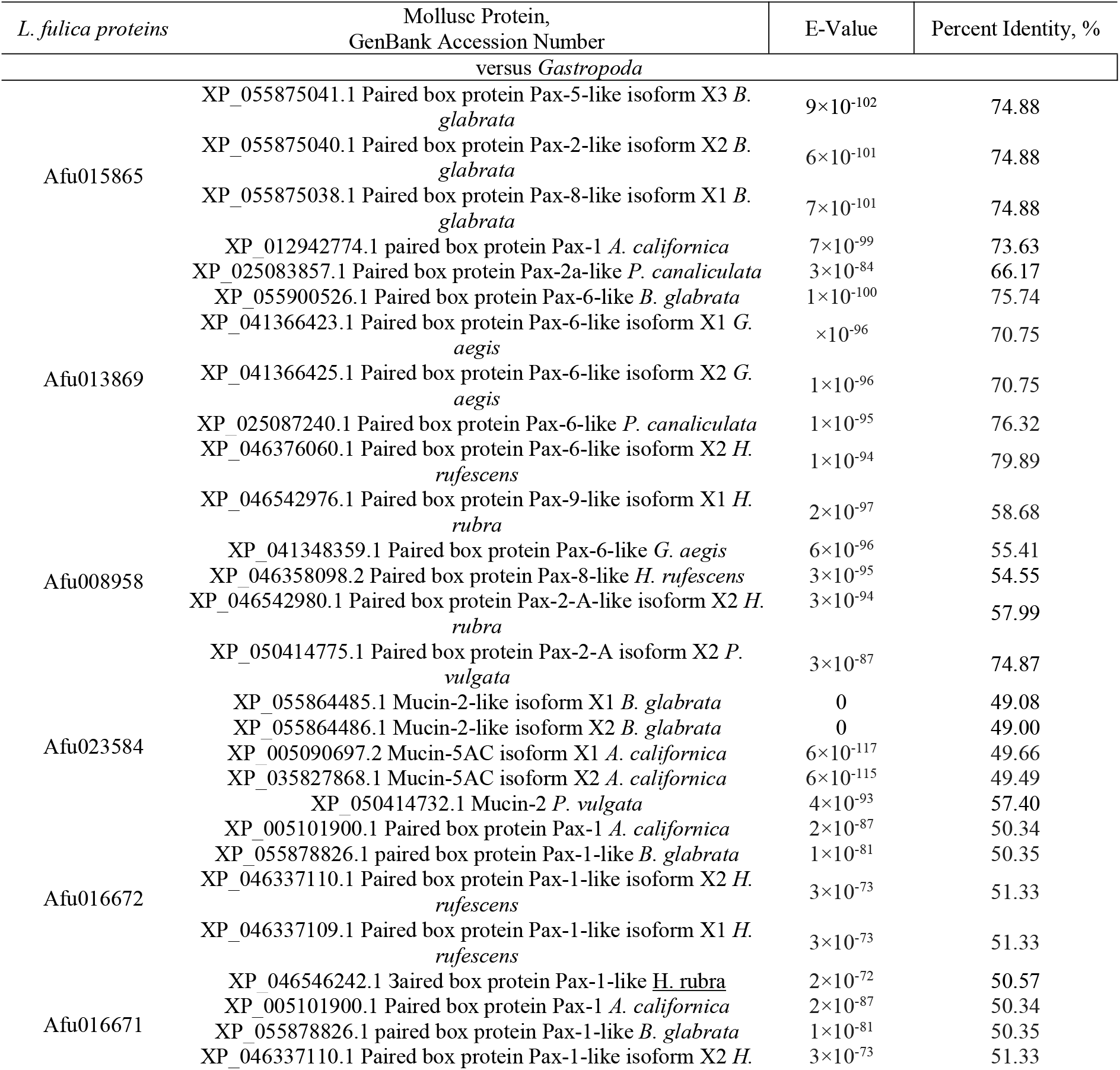
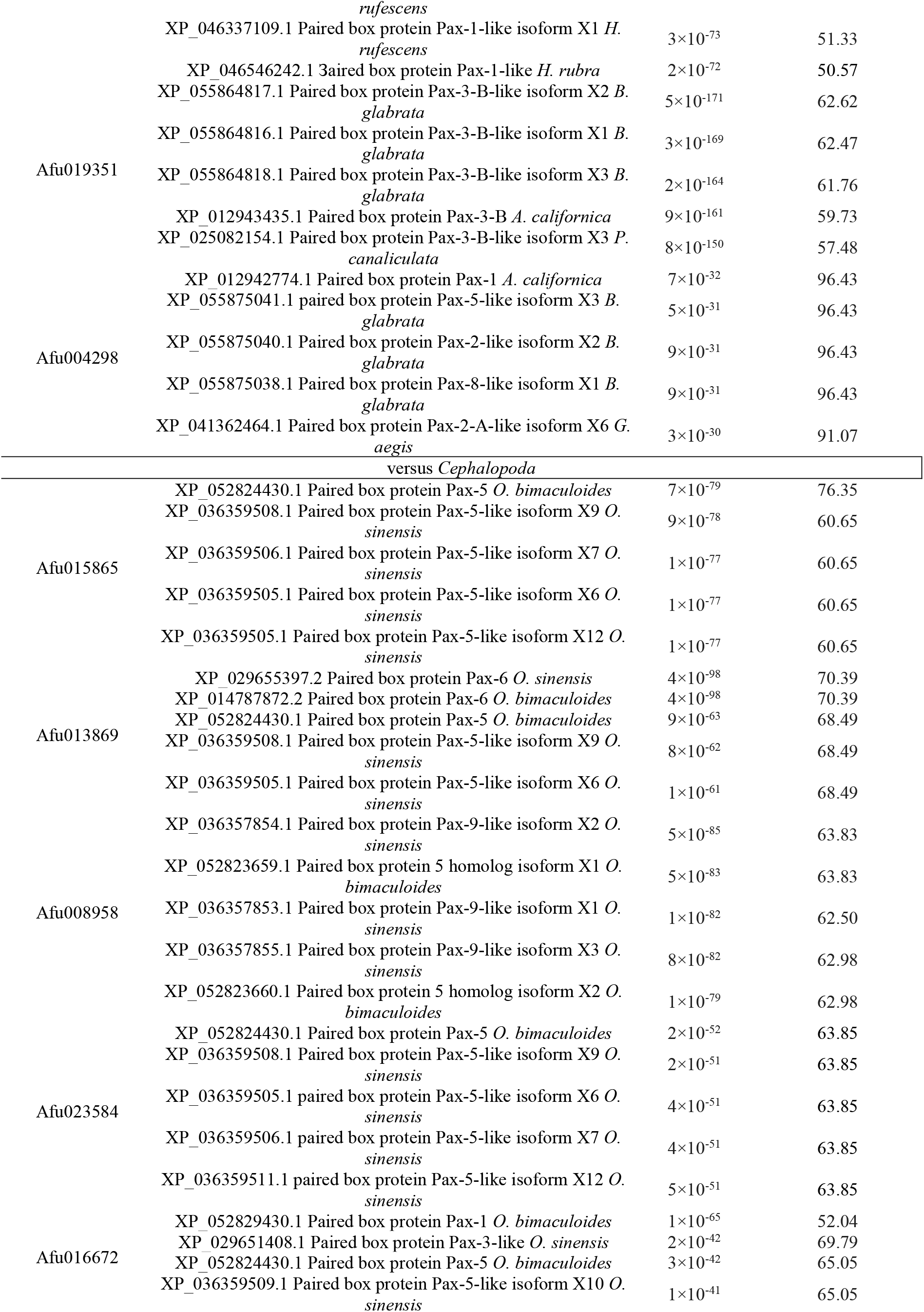
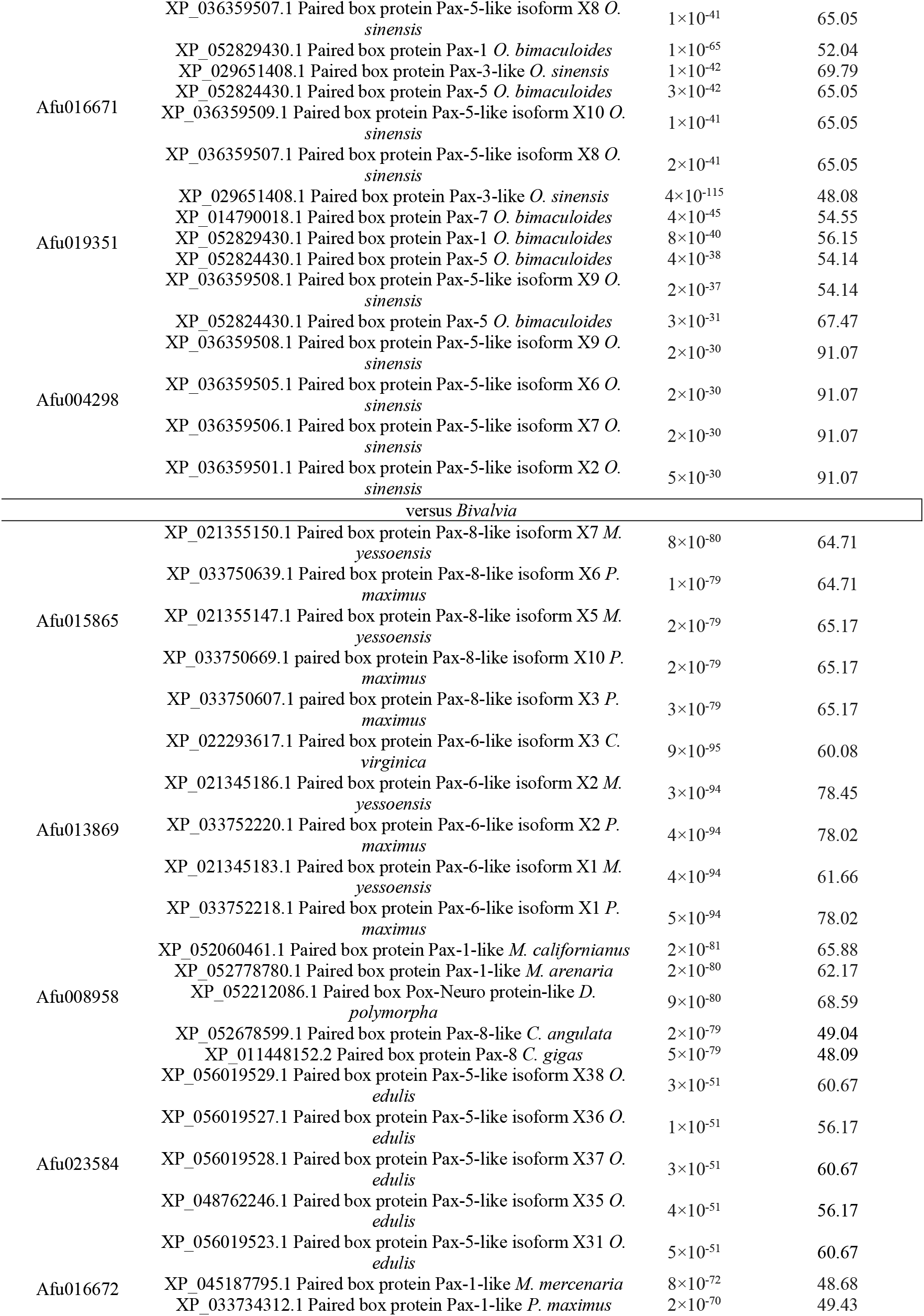
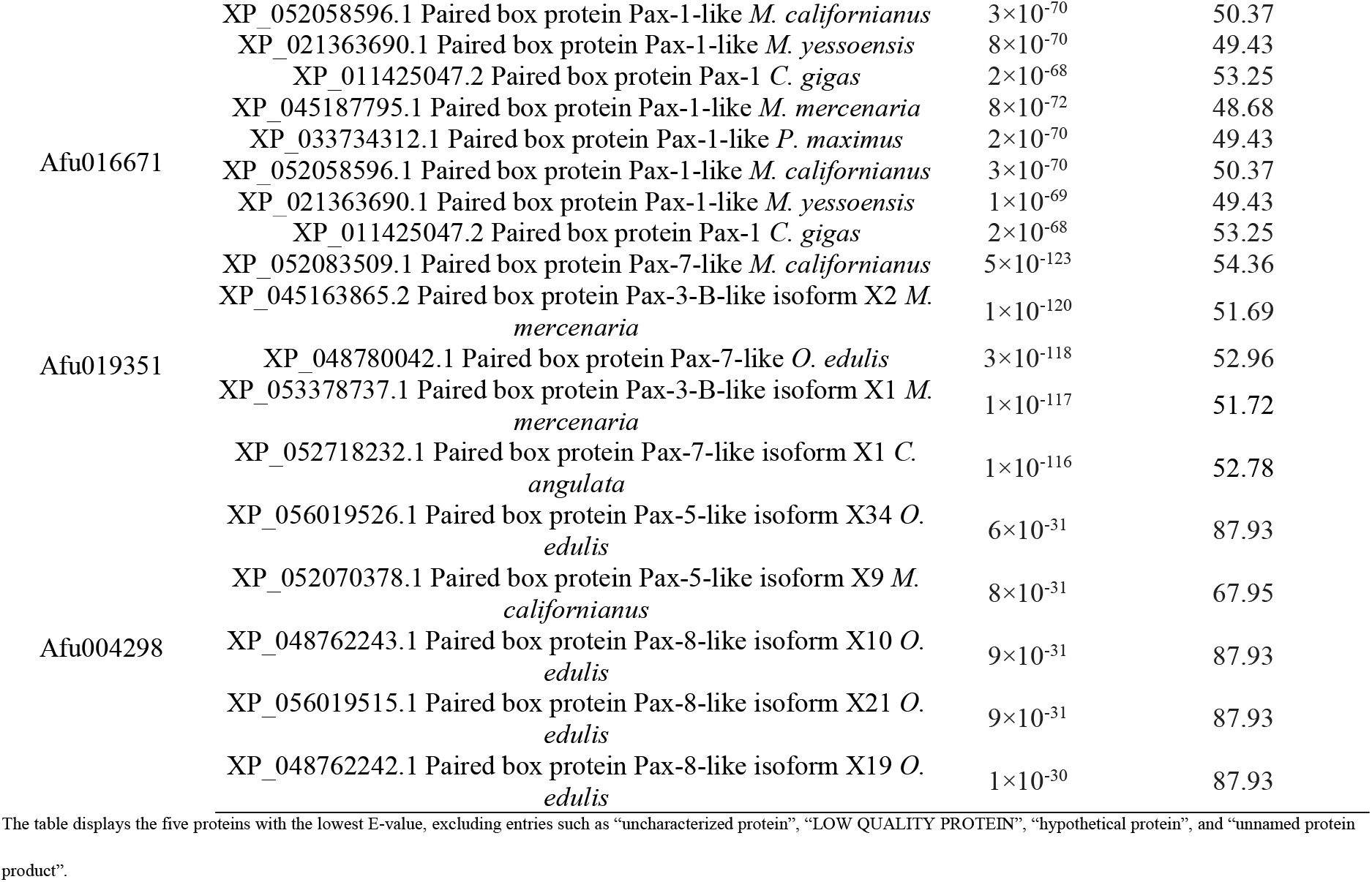
Percent identity of *Lissachatina fulica* paired-box-containing predicted proteins versus *Gastropoda, Cephalopoda* and *Bivalvia* mollusc proteins by BLAST.

Based on the results of identity determination using BLAST (Table 3), it is evident that there are no definitive matches with the findings from phylogenetic analysis and the identification of homeodomain and octapeptide sequences. However, there is some overlap observed across PAX protein subfamilies. For instance, the BLAST analysis revealed the highest similarity of Afu016671 and Afu016672 to PAX-1 proteins (i.e., to subfamily I) found in gastropod and bivalve molluscs. The determination made through both phylogenetic and BLAST analyses for Afu013869 (PAX6) and for Afu019351 (subfamily III) also matched.

### Discussion and conclusions

In this study, we first identified Paired box proteins in *Lissachatina fulica*, and the domain structure of these amino acid sequences is shown in Fig. 4.

**Fig. 4.**
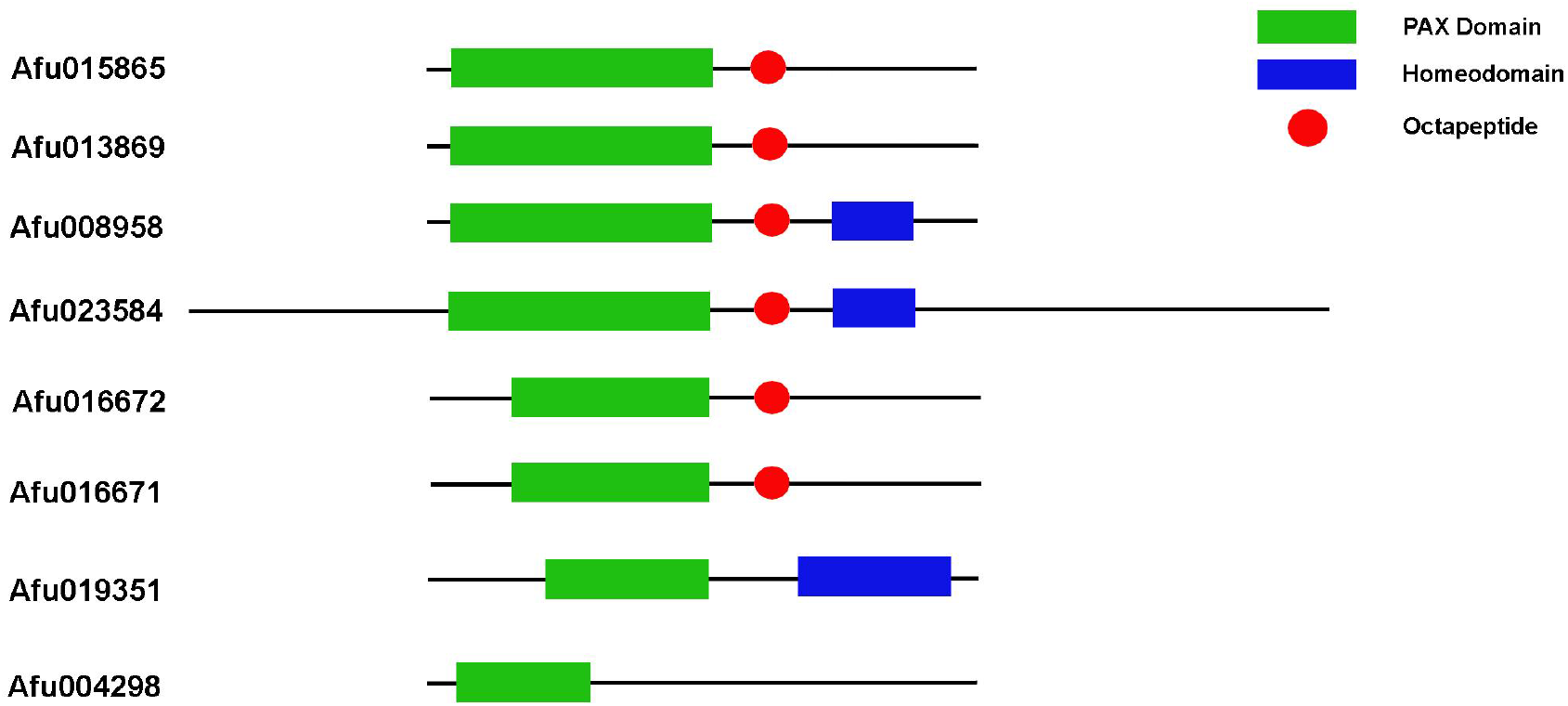
Domain structure of predicted PAX family proteins of *Lissachatina fulica*. The image was created using GNU Image Manipulation Program (GIMP) (version 2.10)

A detailed study of the domain structure, phylogenetic and BLAST analyses allowed us to assign the Afu016671 and Afu016672 sequences to subfamily I of PAX. These proteins contain octapeptides typical for sequences of the subfamily and, conversely, are lack of homeodomain. The deduction is consistent with the data reported by Navet et al. (Navet et al. 2017). Wherein, it should be noted that these sequences contain an incomplete Paired-box domain, which may indicate the similarity of their sequences to Lune/eye gone (Eyg) a PAX-like protein found in *Drosophila*. The Eyg protein of *Drosophila* lacks the N-terminal PAI domain but contains the C-terminal RED domain in complex with the homeodomain that differentiates it from Afu016671 and Afu016672, which contrariwise lack the homeodomain (Jun et al. 1998). At the same time, it is believed that the PAI domain is strictly necessary for binding to DNA, whereas the RED domain plays an auxiliary role and is needed only under certain circumstances, for example, in cases of mutation of both the paired domain and its recognition sequence (Czerny et al. 1993). It should be noted that despite the high homology of these two sequences, there are still structural differences between Afu016671 and Afu016672, which are confirmed by docking data. Afu016672 has lower HADDOCK score (−47.8 ± 8. 5) and z-score (−1.6) compared to Afu016671 (−40.1 ± 3.6 and -1.1, respectively), indicating a higher its affinity to DNA, as well as the result of 3D structure alignment (Fig. 5), which obtained an RMSD of 3.940 Å.

**Fig. 5.**
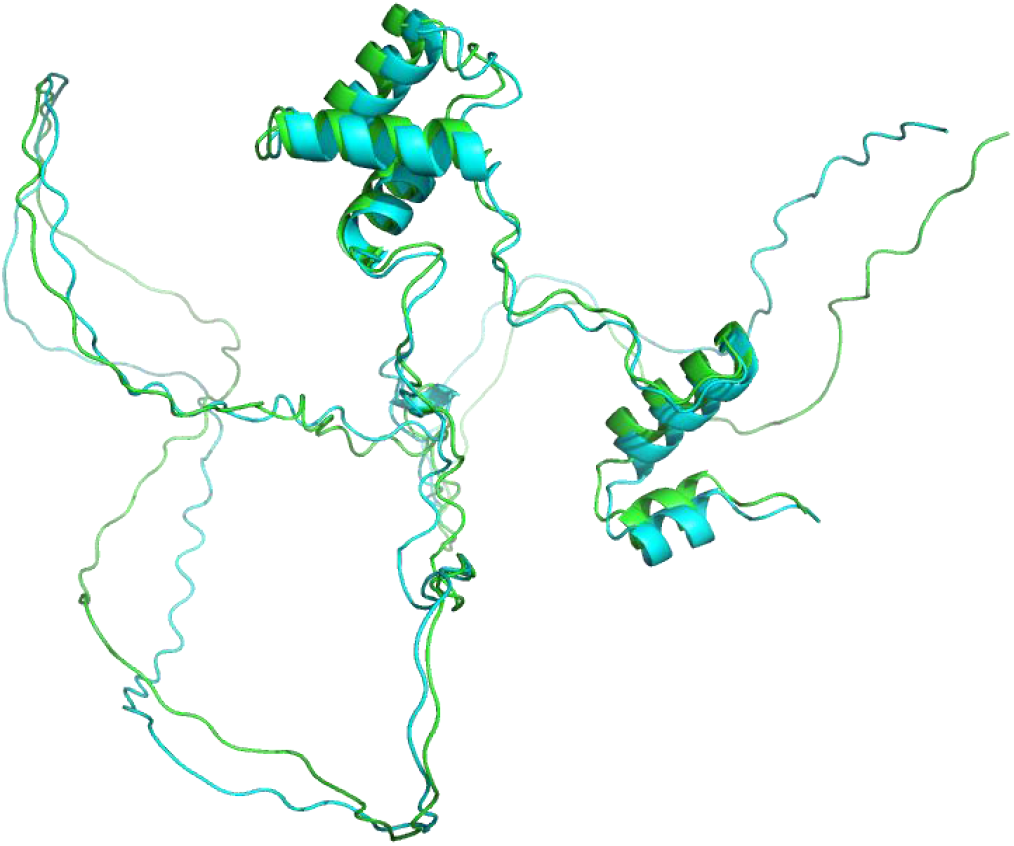
Alignment of 3D structures of the amino acid sequences of Afu016671 (green) with Afu016672 (cyan). The root mean square deviation (RMSD) of the two structures is 3.940 Å. The image was created using PyMOL (version 2.5.7)

The sequences Afu015865 and Afu004298 can also be assigned to the PAX I subfamily. Both of these sequences occupy an intermediate position on the phylogenetic tree between subfamily I and II proteins, forming a common clade with PAX1. At the same time, BLAST analysis almost unambiguously assigns them to subfamily II. However, the absence of a Lys-Arg-rich region at the C-terminus, characteristic of subfamily II PAX proteins (Wollesen et al. 2015), does not allow us to assert Afu015865 and Afu004298 to this subfamily. It is worth noting, wherein, that Afu015865 contains all the necessary components: a complete Paired-box domain and octapeptide (Fig. 4) to be a PAX1 protein. Wherein Afu004298 contains only a PAI domain (Fig. 1), which, as noted above, may be a sufficient condition for this sequence to have DNA-binding properties. The latter is supported by the docking data, as Afu004298 has lower HADDOCK score and z-score values compared to Afu015865: -115.2 ± 18.3 and -2.3 vs. -48.3 ± 3.5 and -1.3, respectively.

A more unambiguous decision about the assignment to the certain PAX family can be made for proteins Afu019351, specifically to subfamily III and, most likely, to PAX3 proteins. Afu019351 and PAX3 proteins of *P. vulgata* form a common clade on the phylogenetic tree, as confirmed by BLAST analysis. Moreover, the structural components (Fig. 4), particularly the presence of a full Paired-box domain and homeodomain, and the absence of an octapeptide indicate membership in subfamily III. It is also worth noting, based on the docking results, the rather strong DNA-binding ability of this protein (HADDOCK score = -42.1 ± 14.6 and z-score = -1.9) compared to the reference Paired-box domain.

The Afu013869 sequence is most likely belongs to the subfamily IV of PAX proteins, as confirmed by phylogenetic and BLAST analyses. However, the domain structure of this protein does not fully correspond to PAX6 proteins, for example, those of cephalopods molluscs, in which these proteins contain a homeodomain in addition to the Paired-box domain. In contrast, Afu013869 does not have a homeodomain but has an octapeptide sequence “YDKLGLLG” that corresponds to PAX subfamily II proteins. However, if we extend the comparison region of this octapeptide, it can be found the slightly modified motif “MYDKLGLLNGQ” (“MYDKLGLLNGH” in *L. fulica*) between the Paired-box domain and the homeodomain. This motif has been shown in the study by Loosli et al. to be a highly conserved structure of PAX-6 proteins found in vertebrates, *Drosophila*, sea urchin, and the nematode *Lineus sanguineus*, but absent in other PAX proteins (Loosli et al. 1996). Furthermore, the C-terminal domain of Afu013869 is represented by a Pro/Ser/Thr-rich region (Yoshida et al. 2014), which is characteristic of some variants of PAX6 proteins in cephalopods, namely squid and cuttlefish. Moreover, the squids can have a large number of PAX6 variants, including classical variants containing the complete Paired-box domain, homeodomain, and Pro/Ser/Thr-rich region, as well as variants with deletions in the homeodomain region or insertions in the Pro/Ser/Thr-rich region and between the Paired-box domain and homeodomain (Yoshida et al. 2014). It’s also shown PAX-6 protein of some molluscs can differ from the canonical variant. For example, *Mytilus galloprovincialis* has two PAX6 isoforms, one of which contains only Paired-box domain, and herewith the same protein PAX6 in *Octopus bimaculoides* lacks the homedomain (Navet et al. 2017). Thus, by annotating Afu013869 as PAX6, we can hypothesize that insertion of the octapeptide sequence and deletion of the homeodomain exons could have occurred without altering the Pro/Ser/Thr-rich region in the *PAX6* gene of *L. fulica*.

Afu023584 can also be assigned, like Afu013869, to a specific PAX protein subfamily. According to all obtained data, except for BLAST analysis, Afu023584 represents a PAX-beta protein. BLAST analysis on gastropod molluscan proteins identified this sequence as Mucin-5AC, which is not uncommon and has already been shown by Navet at al. (Navet et al. 2017). In their study the sequence XP_005090697.1 Mucin-5AC isoform X1 (an earlier version of XP_005090697.2) of *Aplysia californica* was used as the true PAX-beta protein because the mRNA of this gene was originally annotated as Pax-beta.

Based on the results of domain structure search and phylogenetic analysis, the Afu008958 sequence can be annotated as Pox-Neuro. In particular, this sequence contains the “VPGLSYPRLV” motif found in Pox-Neuro proteins of some *Mollusca* species (Navet et al. 2017). However, Afu008958 also contains a partial homeodomain, which is not typical for Pox-Neuro proteins of *Lophotrochozoa*. Similar to PAX6, the variability of this protein can be supposed. Further, a more in-depth and detailed study of Pox-Neuro protein structures is complicated by the restricted number of available annotated sequences, as well as the absence of this PAX protein subfamily in chordates (Matus et al. 2007).

In summary, based on the obtained data, it can be concluded that the *L. fulica* genome contains genes encoding amino acid sequences corresponding to 5 subfamilies of PAX proteins (I, III, IV, β, and Pox-Neuro). It should be noted that the genes encoding PAX proteins are prone to multiple alternative splicing, and as a result, several isoforms may be present simultaneously in the same tissues of the *L. fulica*. In contrast, it is suggested that co-expressed isoforms are evolutionarily more restricted than each individual isoform, as orthologous evolutionarily conserved isoforms may have different transcriptional activities in different species (Short et al. 2012). In particular, the utilization of the PAI, RED, and homeodomain domains by PAX proteins can potentially enable their involvement in diverse developmental processes at different stages. This versatility allows them to regulate specific target genes and contribute to different aspects of development.

Therefore, the further stage of studying the identified sequences for their functional differentiation will involve searching for transcripts of genes encoding these sequences and their corresponding amino acid sequences in tissues of *L. fulica* at different stages of development. This will provide insights into their potential involvement in the mechanisms of morphogenesis and regeneration in this gastropod mollusc.

## Supplementary Information

The online version contains supplementary material available at https://doi.org/XXX.

## Author contributions

Conceptualization: Irina N. Dominova and Valery V. Zhukov; Methodology: Irina N. Dominova; Formal analysis and investigation: Irina N. Dominova, Kristina Golovneva and Nadezhda Korshunova; Data curation, I.N.D.; Writing—original draft preparation: Irina N. Dominova and Valery V. Zhukov; Writing—review and editing: Irina N. Dominova and Valery V. Zhukov; Funding acquisition: Valery V. Zhukov; Resources: Irina N. Dominova; Supervision; Valery V. Zhukov. All authors have read and agreed to the published version of the manuscript.

